# Deciphering conformational preferences of RNA in protein-RNA recognition

**DOI:** 10.64898/2026.05.14.725147

**Authors:** Shri Kant, Savan Masipeddi, Ranjit Prasad Bahadur

## Abstract

Conformational plasticity of RNAs plays important roles in recognizing RNA-binding proteins, and is often modulated by their binding partners. Here, we investigate RNA conformational preferences in a non-redundant dataset of 263 protein-RNA complexes to characterize the structural landscape associated with protein recognition. RNA dinucleotide segments are analyzed using seven backbone torsion angles (δ_1_, ε_1_, ζ_1_, α_2_, β_2_, γ_2_, and δ_2_), two glycosidic torsion angles (χ_1_ and χ_2_) and the pseudo-torsion angle μ. Focusing on dinucleotide steps present in both interface and non-interface regions, we performed density-based clustering using selected backbone torsion angles to identify recurrent conformational states. We identify 28 distinct RNA dinucleotide conformers containing at least ten members each. Among these, eight conformers represent previously unreported nucleotide conformers (NtCs), including the transitional and the non-canonical states AB06, AB07, BB21, BB22, OP32, OP33, IC08 and IC09. Several of these conformers are preferentially enriched at protein-binding interfaces, suggesting their involvement in local conformational adaptation during protein-RNA recognition. The newly identified conformers span transitional A-B geometries, distorted B-like states, open conformations and compact intercalated structures, highlighting the remarkable structural plasticity of RNA in ribonucleoprotein complexes. Overall, this study expands the current understanding of RNA conformational space and provides a refined RNA dinucleotide conformer library for protein-RNA complexes. These findings will facilitate the identification of novel RNA structural motifs and improved RNA structural modeling, docking protein-RNA complexes and deep learning-based prediction frameworks for describing RNA tertiary structures.

## Introduction

RNA molecules are considered as the bridge between DNA and proteins. In recent decades, scientists have recognized its remarkable functional diversity encompassing protein-coding, regulatory and catalytic roles (Hentze et al. 2018; Vicens and Kieft 2022; Liao et al. 2025; Eisenstein 2025). The remarkable structural flexibility of RNA allows it to assume diverse and complex three-dimensional (3D) conformations essential for its function (Hentze et al. 2018; Vicens and Kieft 2022). High-resolution techniques, including X-ray crystallography, provide the atomic intricacies of RNA. Nevertheless, acquiring high-quality crystals is often difficult owing to the inherent flexibility of RNA (Eisenstein 2025). Cryo-electron microscopy (cryo-EM) is a promising alternative for examining RNA structures. It can capture RNA molecules in near-native forms at almost atomic resolution without crystallization (Bonilla and Jang 2024). Kretsch *et al*. (2025) recently determined an RNA-only cryo-EM structure comprising over 800 nucleotides, highlighting the increasing interest in studying 3D structures of RNA at atomic resolution (Kretsch et al. 2025). Although the 3D structures of RNA have increased rapidly in recent years in Protein Data Bank (PDB) (Berman 2000), the extensive structural variation among RNA families remains inadequately documented. Among the 4,178 RNA classes catalogued in the RFAM 15.0 database (Ontiveros-Palacios et al. 2025), merely 143 have their tertiary structures experimentally elucidated. Although the knowledge of 3D structure is essential for understanding RNA function, yet examining secondary structure is a fundamental and pragmatic approach to investigating RNA architecture and its dynamics (Wayment-Steele et al. 2022; Bu et al. 2025).

Traditional representations of RNA structure, such as the A-form helix, fail to capture the rich polymorphic conformations observed in functional RNAs (Bernard et al. 2024; Bonilla et al. 2024; Bu et al. 2025). These include hairpin, junction, pseudoknot, G-quadruplex, triple helices, three- and four-way junction, four-stranded G-quadruplex or parallel helices, and an unusual A-A helix (Leontis and Westhof 2001; Vicens and Kieft 2022; Wayment-Steele et al. 2022). Their existence indicates that RNA structure is much more polymorphic than it might be deduced from the misleading simplicity of the canonical A-RNA (Keating et al. 2011; Richardson et al. 2008; Černý et al. 2020). Foundational work by Altona and Sundaralingam (1972) introduced the concept of pseudorotation to describe sugar conformations (Altona and Sundaralingam 1972). Later efforts, such as the η/θ pseudo-torsional angle formalism (Humphris-Narayanan and Pyle 2012; Keating and Pyle 2012), significantly simplified RNA backbone analysis. The introduction of RNA backbone rotamer libraries (Murray et al. 2003a, 2005), RNA conformational classification (Schneider 2004) and discrete RNA backbone suite conformers (Richardson et al. 2008) enable systematic classification of RNA conformations. More recent framework DNATCO (Černý et al. 2020), Conformational Alphabet of Nucleic Acids (CANA) extends this classification to a broader dinucleotide level, facilitating motif recognition, modelling and RNA design. However, this standard classification of the diNucleotide Conformers (NtCs) has not been explicitly explored in protein-RNA complexes or ribonucleoproteins (RNP). Therefore, a specialized RNA conformer library is essential to understand their role in protein-RNA recognition (Mukherjee and Bahadur 2018; Batey et al. 2001; Kligun and Mandel-Gutfreund 2015).

In this study, we have performed a comprehensive conformational analysis of 6,740 RNA dinucleotide steps derived from a non-redundant benchmark dataset of 263 protein-RNA complexes and 145 RNA-only structures curated from the Protein Data Bank. The refined RNA dinucleotide conformer library distinguishes conformational preferences between protein-RNA interface and non-interface regions, thereby capturing context-dependent RNA structural variability associated with molecular recognition. Our analyses demonstrate that the canonical A-form conformer AA00 remains the predominant RNA structural state. At the same time, several non-canonical and flexible conformers, particularly OP22 and OP25, are significantly enriched in protein-binding regions and are frequently associated with flexible single-stranded RNAs. Importantly, we identified eight dinucleotide conformers AB06, AB07, BB21, BB22, OP32, OP33, IC08, and IC09, which are not reported in earlier studies. These conformers exhibit distinct structural features and preferential enrichment at protein-binding interfaces. We further observed that AA18 is exclusively associated with duplex RNAs, consistent with canonical stacking and base-pairing interactions, whereas AA10 corresponds to Mg²⁺-stabilized intercalated geometry. Collectively, these findings highlight the importance of RNA conformational plasticity in protein recognition and provide a specialized conformational framework that will facilitate RNA structural modeling (Ou et al. 2022), flexible protein-RNA docking (Tuszynska et al. 2014), RNA inverse design (Joshi and Liò 2025) and next-generation RNA structure prediction methods (Townshend et al. 2021; Boyd et al. 2023; Sumi et al. 2024).

## Materials and methods

### Dataset of protein-RNA complexes

In this work, we curated a non-redundant dataset of protein-RNA complexes from the PDB at resolutions of 2.5 Å or better, comprising 263 complexes with a maximum 35% sequence identity (Kant et al. 2025) cutoff. Based on the type of the interacting RNA, the complexes are classified into four structural classes: (A) complexes with tRNA, (B) complexes with rRNA, (C) complexes with duplex RNA, and (D) complexes with single-stranded RNA (Bahadur et al. 2008).

### A non-redundant dataset of free RNA

We have curated a dataset of 145 free RNA structures from the PDB. We identified these RNAs by searching for the keyword “RNA” while excluding proteins, DNA, ligands and hybrid structures solved by Cryo-EM and X-ray crystallography. Each RNA chain in our dataset has at least five nucleotides. We made the dataset non-redundant by restricting sequence identity between any two chains to 35% using BLAST (Boratyn et al. 2013). When multiple entries of the same RNA structure satisfy the above criteria, we selected the one with better resolution, fewer missing nucleotides and fewer gaps. Finally, we examined all the structures for modified or incomplete nucleotides. The PDB files were cleaned, and alternate conformations and occupancy were selected using pdb-tools (Rodrigues et al. 2018).

### Identification of interface residue and nucleotides and estimation of interface area

NACCESS (Hubbard and Thornton 1993) was used to identify the interface area between protein and RNA with a default probe radius of 1.4 Å. We identified residues and nucleotides at the protein-RNA interface by estimating their solvent-accessible surface area (SASA). The interface area is the sum of the SASA of protein and RNA less that of their complex. All atoms of residues and nucleotides that lose SASA upon complexation are considered at the interface.

### Calculation of protein-RNA interactions

Hydrogen bond (H-bond) interactions between proteins and RNA molecules were identified using the HBPLUS program with default geometric criteria (McDonald and Thornton 1994). HBPLUS detects H-bonds based on donor-acceptor distance and angular constraints by incorporating both explicit and inferred hydrogen atoms. Base-stacking and protein-RNA stacking interactions were calculated using an in-house Python script. A stacking interaction was defined when the centroid-centroid distance between the aromatic ring was ≤ 4.5 Å and the angle between the ring planes was ≤ 35° (Barik et al. 2016).

### Defining geometric parameters of RNA

We quantified the following geometric parameters of each dinucleotide: seven backbone torsion angles (δ_1_, ε_1_, ζ_1_, α_2_, β_2_, γ_2_ and δ_2_), two torsions around the glycosidic bonds (χ_1_ and χ_2_), a pseudo-torsion angle (μ) and two distance measurements (NN and C′C′). For each dinucleotide, the atoms involved in calculating the backbone dihedral angles are: C5′(1)-C4′(1)-C3′(1)-O3′(1) for δ_1_, C4′(1)-C3′(1)-O3′(1)-P(2) for ε_1_, C3′(1)-O3′(1)-P(2)-O5′(2) for ζ_1_, O3′(1)-P(2)-O5′(2)-C5′(2) for α_2_, P(2)-O5′(2)-C5′(2)-C4′(2) for β_2_, O5′(2)-C5′(2)-C4′(2)-C3′(2) for γ_2_ and C5′(2)-C4′(2)-C3′(2)-O3′(2) for δ_2_ (Figure 1).

**Figure 1.**
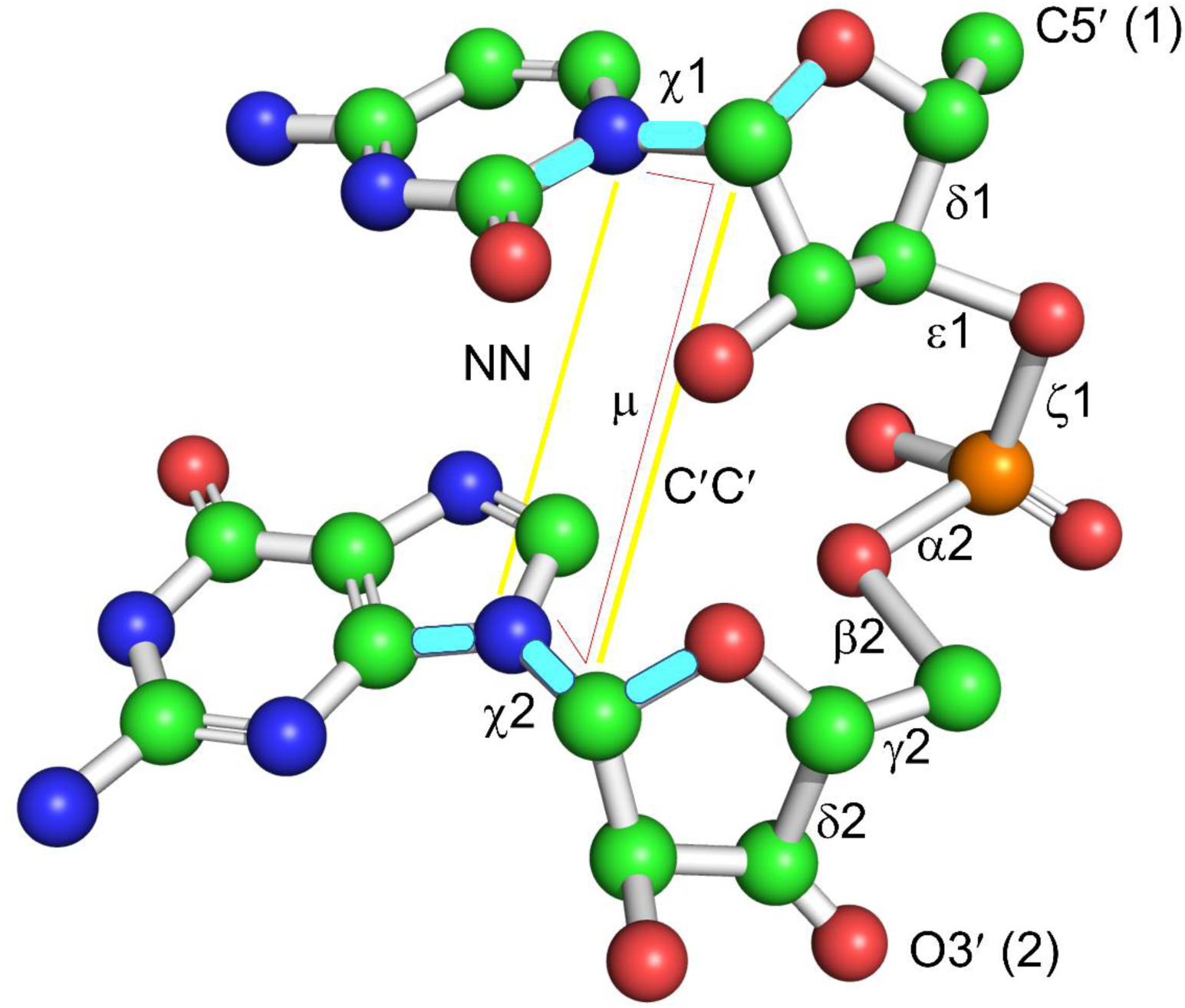
Schematic representation of the seven backbone torsion angles δ_1_, ε_1_, ζ_1_, α_2_, β_2_, γ_2_, δ_2_, and two side-chain torsions around the glycosidic bonds χ_1_ and χ_2_. Additionally, a pseudo-torsion (μ) is defined as the torsion between atoms defining the glycosidic bonds of the first and second nucleotide N1/N9(1)-C1′(1)-C1′(2)-N1/N9(2) (shown in red line). The NN distance is between side-chain N1/9 and N9/1 atoms. The C′C′ distance is between C′1(1)-C′1(2) (shown in yellow lines).

The side-chain orientation was captured by the glycosidic χ angle including the following atoms: O4′(1)-C1′(1)-N1/N9(1)-C2/4(1) for χ_1_ and O4′(2)-C1′(2)-N1/N9(2)-C2/4(2) for χ_2_. The pseudo-torsion angle (μ) represents the torsion between the atoms defining the glycosidic bonds of the first and the second nucleotides N1/N9(1)-C1′(1)-C1′(2)-N1/N9(2). We also included two distance parameters: NN is the distance between N1/N9 of the first nucleotide and N1/N9 of the second nucleotide, and C′C′ is the distance between C1′ of the first nucleotide and C1′ of the second nucleotide (Černý et al. 2020). Additionally, we have calculated five internal sugar torsions, v0 to v4, which describe the puckering of the ribose ring (Altona and Sundaralingam 1972). Together, these backbone, sugar and glycosidic angles provide a comprehensive geometric description of RNA conformational space, forming the basis for all subsequent analyses.

### Construction of dinucleotide RNA conformer libraries

Identifying new conformers from a large and heterogeneous conformational space requires a clustering approach that navigates two competing size constraints: clusters that are too small risk lacking enough signal to confidently identify conformers, while clusters that are too large risk diluting the signal from individual conformers. We applied Density-Based Spatial Clustering of Applications with Noise (DBSCAN) (Ester et al. 1996) to cluster backbone torsional angles, choosing it for its ability to identify clusters of arbitrary shapes. DBSCAN identifies clusters by locating core density regions, where at least *MinPts* points fall within a radius ε (epsilon). Points within ε distance of a core point are included in the cluster as border points, while points that satisfy neither condition are labelled as noise. To handle the heterogeneous distribution of the torsional angles, an iterative approach was used where DBSCAN was applied in multiple successive stages with adaptive parameters (Figure S1). All the points unassigned to any cluster are labelled as outliers. Visualizing and analyzing higher-dimensional data is not feasible since it is impossible to represent seven-dimensional data in two or three dimensions. To overcome this problem, we used Uniform Manifold Approximation and Projection (UMAP) to reduce the dimensionality in torsional space. The parameter “n_neighbors” determines the number of neighbors considered for local approximations, influencing the balance between local and global structures. We set “min_dist” to 0.15, which controls the minimum distance between embedded points, effectively impacting the clustering and distribution of points in lower dimensions. To define the desired dimensionality of the output space, we specified “n_components” to 2, which resulted in a clear and concise 2D representation. Through careful parameter tuning, UMAP facilitates the extraction of meaningful patterns and relationships within the torsional space, enhancing the effectiveness of subsequent analyses and interpretations.

### Assignment of the NtC and CANA

The Confal score is a quantitative metric used to evaluate how well a set of observed torsion angles in a dinucleotide conformer matches a reference conformer class. It is based on the z-score distance of each torsion angle from its corresponding mean in the standard reference set, normalized by the expected standard deviation. Transformed Confal score (scaled from 0 to 100) is given by:

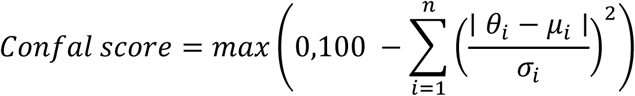

θ_i_: Observed torsion angle *θ_i_* ∈ [0°, 360°]

μ_i_: Cluster mean of torsion angles of the reference conformer class *μ*_*i*_ ∈ [0^°^, 360^°^]

σ_i_: Standard deviation of torsion angles in the reference class

n: Total number of angles

## Results

We have analyzed 6,740 dinucleotides from 263 non-redundant protein-RNA complexes. To understand the conformational changes that enable RNA to adopt various conformational states, we examined 12 geometric parameters of RNA illustrated in Figure 1 (described in the Materials and Methods section). Further, we have categorized the dinucleotide steps based on the location of the i^th^ or the (i+1)^th^ nucleotide present at the interface. This classification yielded 4,266 interface dinucleotides (63.29% of the total dinucleotides present in the dataset) and 2,474 non-interface dinucleotides (36.7%). Additionally, we have also calculated 12 geometric parameters (described in Materials and Methods section) for 12,686 dinucleotides from a non-redundant dataset of 145 free RNA structures.

### Distribution of the torsion angles

The distribution of each of the six backbone torsional angles, the glycosidic torsional angle and the pseudo glycosidic torsion angle in bound and free RNA is shown in Supplementary Figure S2. Torsion angles δ_1_ show a bimodal distribution with a sharp peak around 84° and a smaller peak around 147° (Figure S2A), which correspond to C3′ endo and C2′ endo ribose puckers, respectively. Torsion angle ε1 shows a skewed Gaussian peak around 200° (Figure S2B). Torsion angle ζ_1_ exhibits a broad distribution with a sharp peak around 286° (Figure S2C). Torsion angle α_2_ shows a trimodal distribution with a sharp peak around 300° along with two smaller peaks around 60° and 160° (Figure S2D). Torsion angle β_2_ shows a Gaussian distribution centered around 180° (Figure S2E). Torsion angle γ_2_ also shows a trimodal distribution with a sharp peak around 60° along with two smaller peaks around 180° and 300° (Figure S2F). A sharp peak in χ₁ around 200° corresponds to the predominant anti-conformation of the nucleotide bases. In addition, a smaller and broader peak near 60° indicates the presence of minor syn conformations. Most χ values cluster between 180° and 240°, with very few occurrences at higher angles, apart from the minor syn population (Figure S2G). The pseudo-glycosidic torsion angle (μ) shows a broader distribution, predominantly ranging from 10° to 60° (Figure S2H). The distributions of all these torsional angles in free-RNA closely resemble to those observed in bound RNA.

Two-dimensional distributions reveal several intriguing interdependencies among backbone torsion angles (γ₂ vs δ₁ in Figure S3A and β₂ vs ε₁ in Figure S3B). The most noteworthy pattern is observed for the phosphodiester bonds, where distinct clusters indicate preferred conformational states signifying plausible coupling between these angles. Understanding the distribution of the glycosidic torsion angle (χ) is crucial for exploring potential relationships between base-pairing pattern and backbone conformation. Notably, significant clustering is evident in the scatterplots involving glycosidic bonds δ₁ vs χ₁ (Figure S3C) and γ₂ vs χ₁ (Figure S3D), highlighting the conformational preferences and the effect of backbone geometry on base orientation. The distribution of these torsional angles is in close agreement with those found in X-PLOR (Parkinson et al. 1996) and CNS (Brünger et al. 1998).

### Clustering of dinucleotide conformations

DBSCAN is applied to cluster all dinucleotide conformations based on the backbone torsional angles (Figure 1). Given the expected heterogeneity of the conformational space, iterative clustering was employed to capture the progressively rare conformations (Figure S1). The first iteration accounted for ∼58% of all the dinucleotides (3,857), while subsequent iterations recovered an additional 1909 dinucleotides distributed across 27 smaller clusters, each containing at least 10 dinucleotides. The mean and standard deviation of backbone and side-chain torsional angles for all the 28 clusters are provided in Table 1.

Additionally, the “-1” cluster contains the unassigned dinucleotides (NANT). These NtCs are specific to bound RNA. A few of them overlap with the previously reported unified conformers for DNA and RNA (Černý et al. 2020). Unlike previous studies, our analysis is based on a non-redundant dataset of protein-RNA complexes, which primarily consists of small to medium-sized RNAs as we excluded large RNP complexes such as ribosomes and viral capsids. Nevertheless, we ensured that our sampling of local geometric parameters encompasses a diverse range of RNA types. A detailed conformational analysis of RNA conformers reveals the structural diversity of backbone geometries, sugar puckers, glycosidic torsions and inter-base arrangements across multiple conformer clusters. Each of them represents a distinct structural conformation specific for recognizing RBPs. The representative NtCs present at the interface (Table 2) and at the non-interface (Supplementary Table S1) of protein-RNA complexes, which are the nearest to each cluster center.

### Dinucleotide conformers in protein-RNA complexes

To systematically classify RNA conformations in protein-RNA complexes, we adopted the established nomenclature from DNATCO, which classifies nucleic acid geometry using CANA and a more detailed NtC scheme. CANA provides a coarse-grained classification using three-letter codes (e.g., AAA, BBB, ZZZ), whereas NtC defines fine-grained structural states using four-letter identifiers (e.g., AA00, BB00, ZZ01) (complete list in Table 2). Some of the conformers identified in this study closely correspond to previously reported classes, particularly AA00, AA01, AA07, AA10 and AA11, which represent variants of A-form helical geometries with distinct backbone torsional signatures (Figure 2, A-E). While these conformers broadly align with earlier structural definition (Černý et al. 2020), our analysis reveals measurable deviations in mean torsion angles and their distributions, highlighting intrinsic conformational variability within each NtC class.

**Figure 2.**
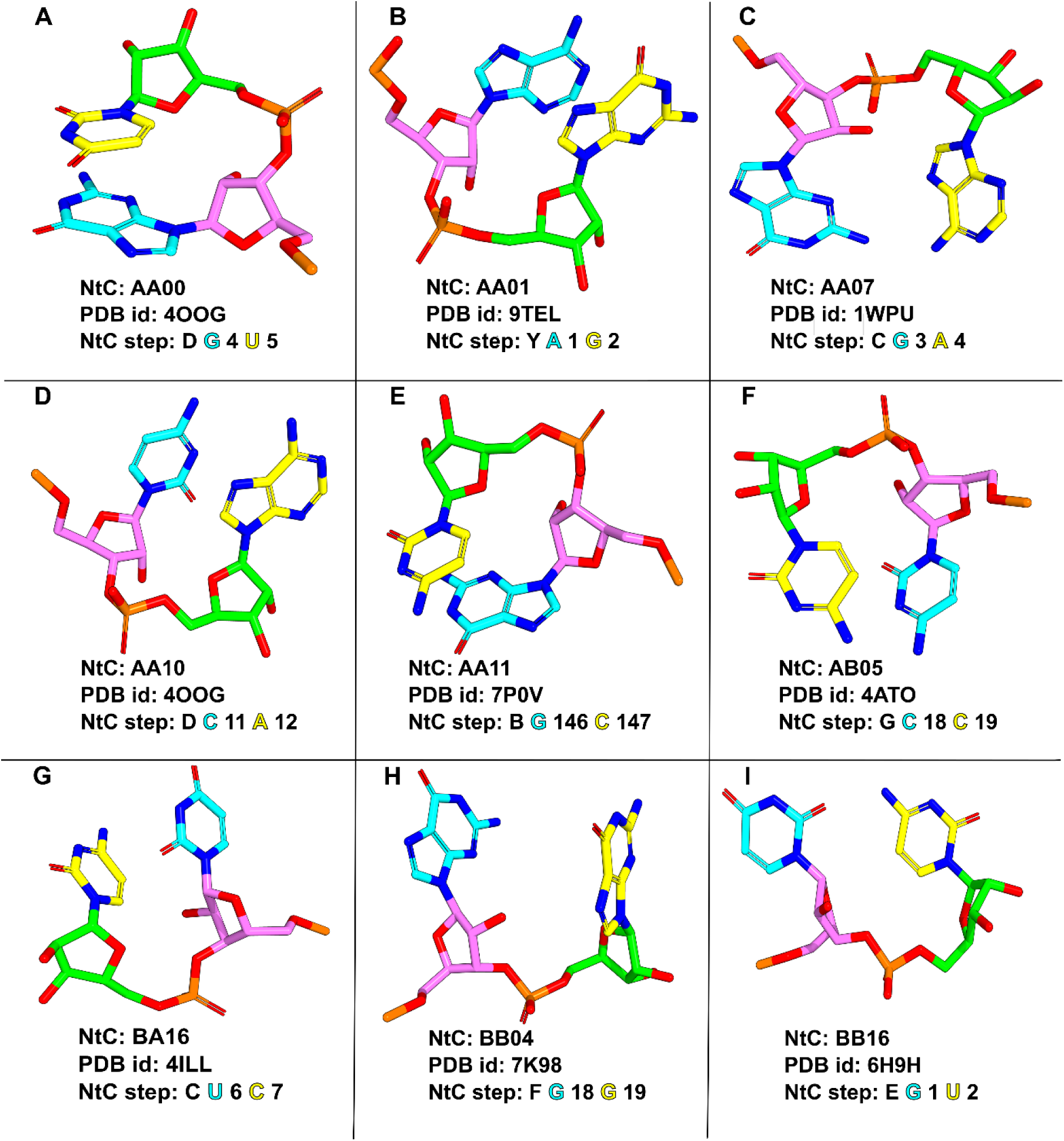
Representative of dinucleotide conformers (NtC) in protein-RNA complexes (A to I). Each panel displays the NtC, PDB ID, and NtC steps along with their chain identifier. The NtC step of the first and the second nucleotide is shown in cyan and yellow colours, respectively.

The canonical AA00 conformer (Figure 2A) represents a well-defined A-form RNA geometry characterized by C3′-endo sugar puckers and anti-glycosidic orientations. In contrast, AA01 (Figure 2B) corresponds to a widened A-like geometry (AAw), maintaining C3′-endo puckers but exhibiting subtle expansion of the helical framework. The AA07 conformer (Figure 2C) represents an unwound A-like state (AAu), characterized by increased base separation and altered backbone torsions. The AA10 (Figure 2D) and the AA11 (Figure 2E) conformers are distinguished by α/γ (80°/220°) torsional switches, where the AA10 often forms intercalated structures stabilized by divalent ions and the AA11 exhibits reduced μ torsion, indicative of enhanced base rotation. Collectively, these conformers expand the structural repertoire of A-form RNA and underscore its adaptability in diverse structural contexts.

In addition to canonical A-like conformations, we also observe transitional states bridging A- and B-form geometries. The A-B class, exemplified by the AB05 (Figure 2F), exhibits mixed sugar puckers (C3′-endo/C2′-endo) and intermediate glycosidic torsions, reflecting a gradual shift from A-like to B-like conformations.

Conversely, the B-A class, represented by the BA16 (Figure 2G), displays reverse transitions with mixed puckers (C2′-endo/C3′-endo) and distinct backbone torsions, including a notable low γ (50°) angle. Furthermore, B-form-related conformers (Figure 2H and 2I) such as the BB04 (B12) and the BB16 capture transitions between BI and BII states or incorporate A-like torsional features within an overall B-like framework, illustrating the structural continuum between canonical helical forms. We have also identified A diverse set of open (OPN) conformers characterized by disrupted base stacking and increased backbone flexibility (Figure 3). These conformers typically exhibit atypical torsion angles and variable sugar puckers, often associated with unstacked or spatially distant bases (Figure 3). However, each conformer has its own unique characteristic. The OP11 (Figure 3A) adopts an open conformation with both the sugars in C3′-endo conformation. It has an unusual ε_1_, ζ_1_, α_2_ and β_2_ torsions, and a low-rise with a mean μ of 100°. Further, the OP11 is characterized by B-A transitional pucker (C2′-endo to C3′-endo) with distant perpendicular bases. On the other hand, the OP15 (Figure 3B) is characterized by B-A transitional pucker with very low γ_2_ (40°) and paired bases. The OP17 (Figure 3C) also exhibits a B-A pucker with a subtle shift in the backbone torsion values. The OP21 (Figure 3D) features a B pucker (C2-endo, C2′-endo) with high μ torsion (220°). The OP22 (Figure 3E) features a B pucker with very low μ torsion (40°), while the OP25 (Figure 3F) exhibits a B-A open conformer with perpendicular bases and very high μ (250°). The OP26 (Figure 3G) exhibits a B-A open conformer with perpendicular bases and ultra-high μ (300°). The OP29 (Figure 3H) exhibits an A-like open conformer with perpendicular bases and μ (120°). Individual OPN subclasses including OP11, OP15, OP17, OP21, OP22, OP25, OP26 and OP29 display distinct combinations of torsional deviations and base orientations, reflecting a spectrum of partially unfolded or dynamically accessible states. Such conformations are frequently enriched in protein-binding regions and may facilitate conformational transitions during molecular recognition.

**Figure 3.**
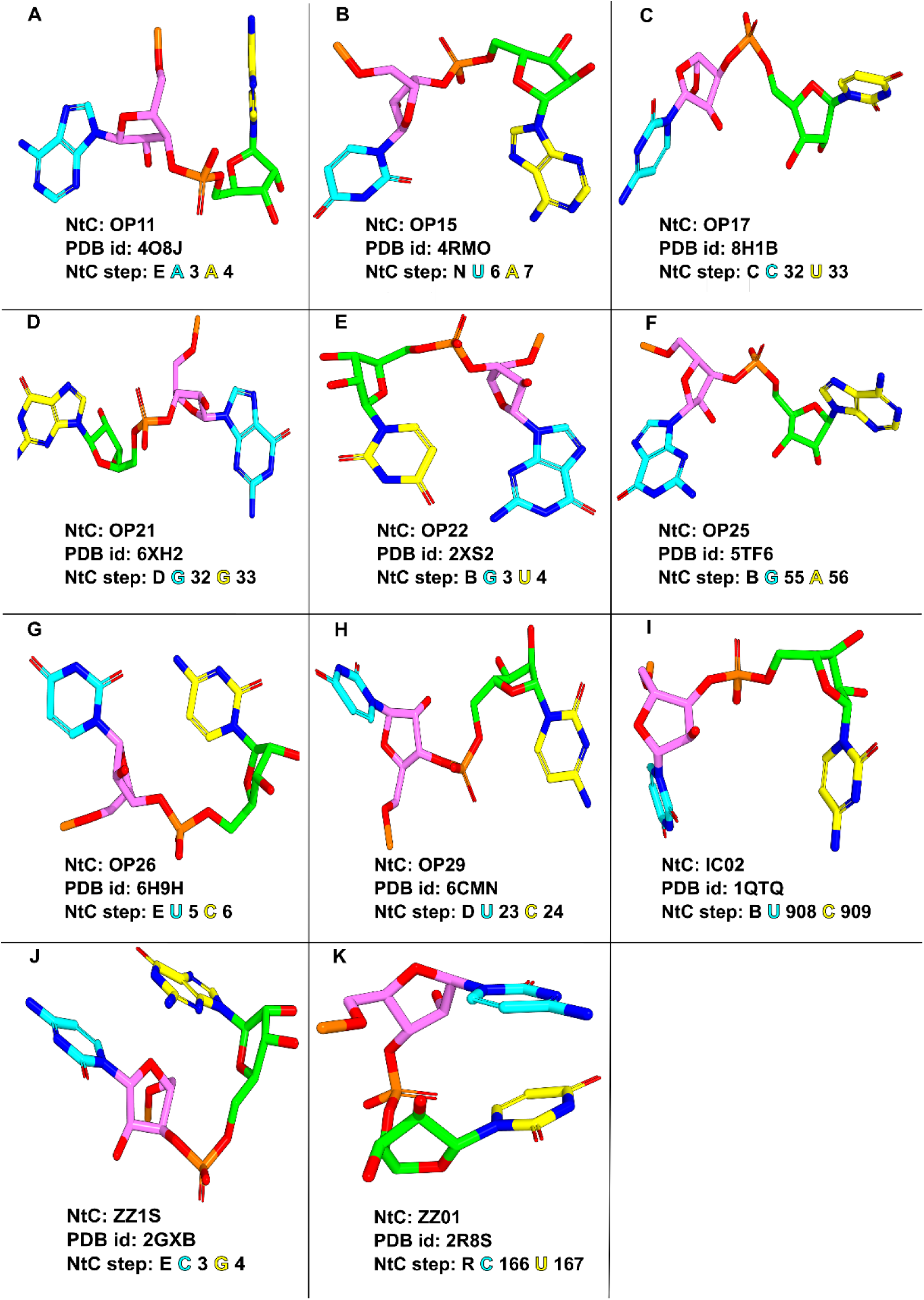
Representative of dinucleotide conformers (NtC) in protein-RNA complexes (A to K). Each panel displays the NtC, PDB ID, and NtC steps along with their chain identifier. The NtC step of the first and the second nucleotide is shown in cyan and yellow colours, respectively.

The intercalated class (ICL), represented by the IC02, exhibits compact geometries with uniform C3′-endo puckers, elevated μ (285°) torsion and characteristic backbone arrangements conducive to stacking interactions. These conformers likely play roles in stabilizing compact RNA motifs or mediating specific intermolecular contacts (Figure 3I). These findings illustrate the rich structural diversity of RNA backbone conformations, many of which are sequence-and context-specific, underpinning RNA’s functional versatility. Finally, Z-like conformers, including ZZ1S and ZZ01, are classified under the ZZZ CANA family (Figure 3J and 3K). These conformers display hallmark features of left-handed helical geometry, such as alternate sugar puckers and syn/anti-glycosidic conformations. ZZ1S, in particular, reflects pyrimidine-purine steps characteristic of Z-RNA. Whereas, ZZ01 represents a Z-like helical variant with distinct α/γ (170°/40°) torsional arrangements, with both the bases adopting the anti-orientation. These conformations may be functionally relevant in regions undergoing structural transitions or adopting non-canonical helical forms.

### Newly reported NtCs from protein-RNA complexes

In this section, we describe a set of newly identified NtCs (Table 1 and Figure 4) derived from our non-redundant dataset of protein-RNA complexes. Notably, these NtCs are not reported in previous classifications, thereby expanding the current conformational landscape captured by the DNATCO framework. We also illustrate the electron density fit of the newly reported NtCs (Figure 5).

**Figure 4.**
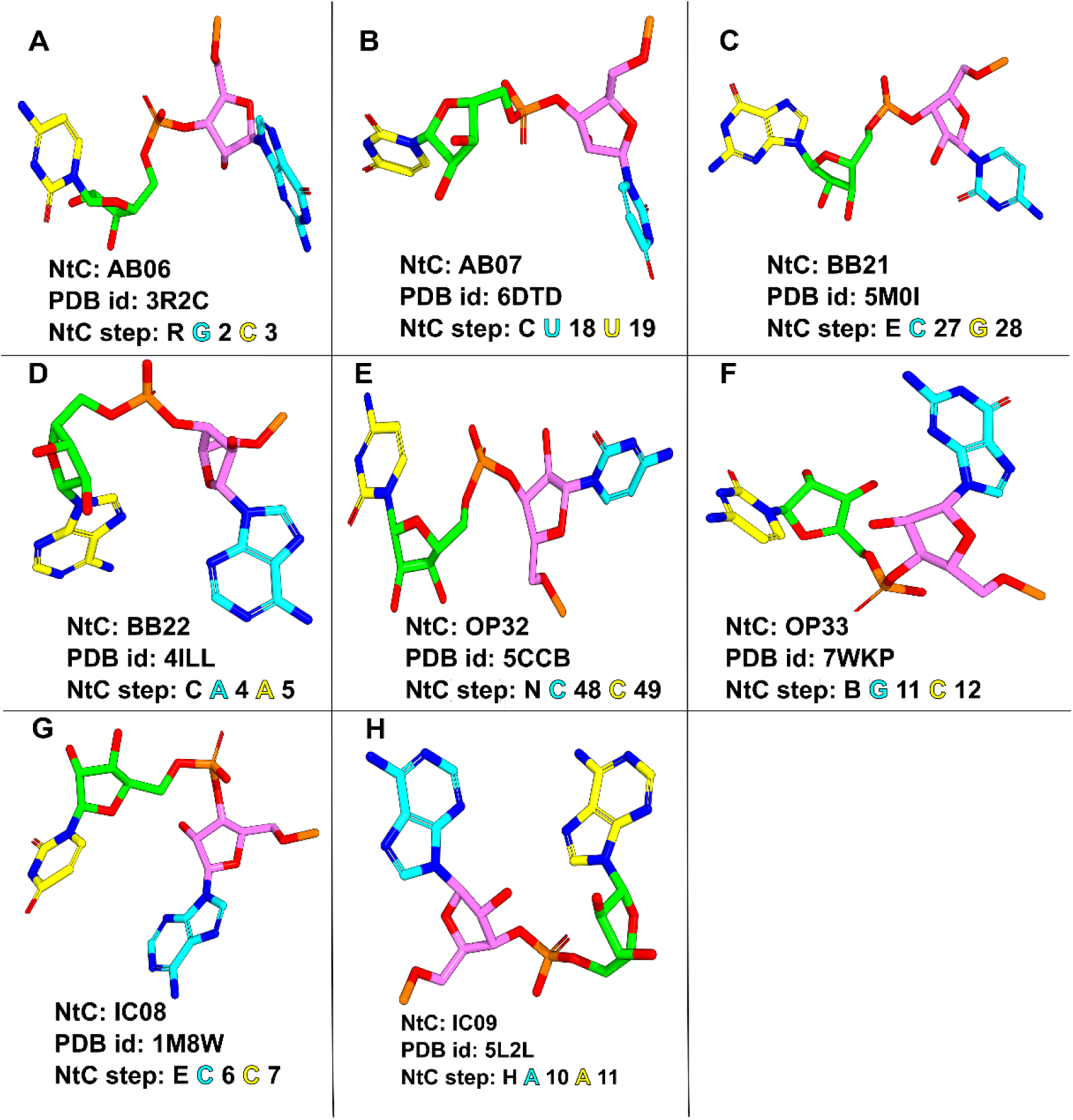
Representative of all newly identified dinucleotide conformers (NtC) in protein-RNA complexes (A to H). Each panel displays the NtC, PDB ID, and NtC steps along with their chain identifier. The NtC step of the first and the second nucleotide is shown in cyan and yellow colours, respectively.

**Figure 5.**
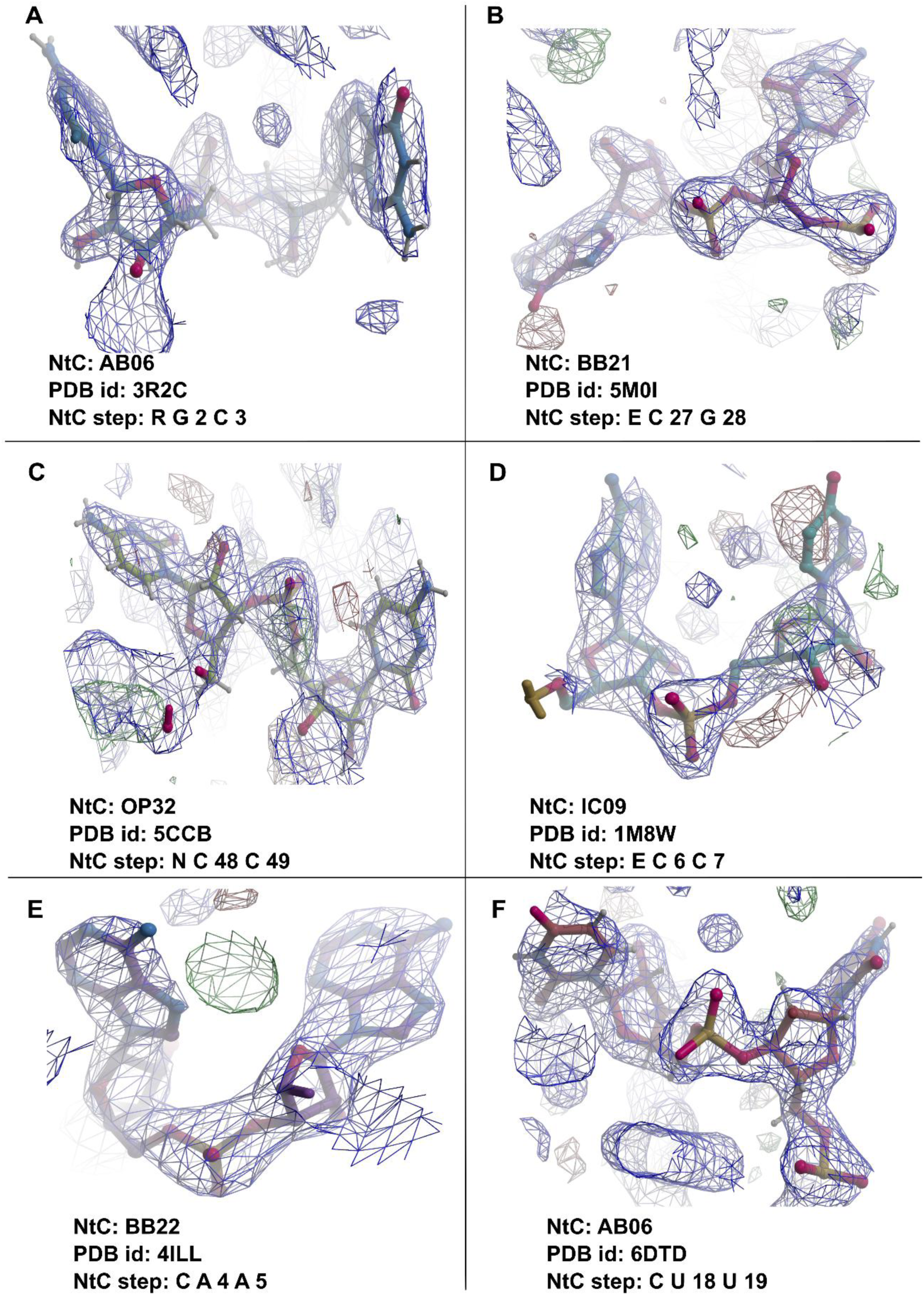
Illustration of the electron density fit of the newly reported NtC steps is shown using Coot software. For each figure NtCs names with PDB ID and dinucleotide steps along with their chain identifier.

The NtCs AB06 and AB07 (Figure 4A and 4B) belong to the A-B transitional CANA class, which represents conformational intermediates between canonical A-form and B-form geometries. In AB06, the dinucleotide adopts a mixed sugar pucker configuration (C3′-endo/C2′-endo) with anti-glycosidic orientations and a relatively low μ torsion (∼70°), indicative of partial unwinding (Figure 4A). In contrast, AB07 (Figure 4B) retains a similar pucker arrangement but exhibits a distinct torsional profile, including reduced α angle and differential glycosidic behavior (χ₁ A-like and χ₂ B-like), accompanied by an elevated μ (∼250°). Both conformers display increased inter-sugar (C1′-C1′) and inter-base (N-N) distances (> 9 Å), placing them at the boundary between the stacked and the open conformations. These features suggest a structurally relaxed state that may facilitate conformational transitions during protein binding.

The conformers BB21 and BB22 (Figure 4C and 4D) are classified within the miB (minor B-like) CANA class, which represents a distorted B-form geometry. BB21 (Figure 4C) exhibits a mixed sugar pucker (C3′-endo/C2′-endo) with a characteristic ε/α torsional switch (high ε ∼260°, low α ∼60°), while maintaining anti-glycosidic orientations and an intermediate μ (∼190°). In contrast, BB22 (Figure 4D) adopts canonical B-type puckers (C2′-endo/C2′-endo) but displays a ζ/α torsional switch (low ζ ∼145°, low α ∼130°) and a marked high μ (∼280°), reflecting significant backbone rearrangement. Both conformers exhibit moderate to large inter-base separations (∼10 Å), indicating partial disruption of stacking. These distorted B-like states likely arise from local structural constraints imposed by protein interactions. The NtCs OP32 and OP33 (Figure 4E and 4F) belong to the open (OPN) CANA class, which is characterized by reduced base stacking and enhanced backbone flexibility. OP32 (Figure 4E) displays a C2′-endo/C3′-endo pucker combination with B-A-like glycosidic orientation (χ₁ ∼275°, χ₂ ∼174°) and is notably enriched at specific positions in folded RNA, where it contributes to tertiary interactions such as cis-Watson-Crick pairing. OP33 (Figure 4F), another open conformer, exhibits a B-A-like pucker arrangement with perpendicular base orientation and a high μ (∼240°), consistent with unstacked geometry. This conformer is also frequently observed at positions mostly folded RNA, known for structural plasticity. The recurrence of these conformers at functionally important sites suggests their role in facilitating RNA folding and tertiary contact formation.

The intercalated conformers IC08 and IC09 (Figure 4G and 4H) are assigned to the ICL CANA class, which represents a compact yet non-helical state. Both the conformers exhibit B-A mixed sugar puckers and pronounced backbone distortions. IC08 (Figure 4G) is characterized by unusually low ζ, α and γ torsions, whereas IC09 (Figure 4H) displays a ζ/α switch (ζ ∼280°, α ∼170°), with both the conformers maintaining B-like anti-glycosidic orientations and high μ values (>310°). The relatively short C1′-C1′ and N-N distances (∼7 Å) indicate tightly packed geometries. Notably, IC09 is predominantly observed at protein-RNA interfaces, where it participates in stabilizing interactions, including H-bonds with protein residues. The moderate B-factor range (18 Å² to 40 Å²) further suggests that these conformations are structurally stable despite their non-canonical nature.

### RNA dinucleotide steps and their preferred conformations in protein-RNA recognition

A standardized Pearson residual (SPR) analysis reveals distinct conformational enrichment patterns (residuals ≥ ±2.0) for RNA dinucleotide steps across interface and non-interface regions of protein-RNA complexes (Supplementary Table S2 and S3). SPR for the free RNA dinucleotides are provided in (Supplementary Table S4). These analyses demonstrate that specific dinucleotide sequences preferentially adopt distinct NtC conformations depending on their structural and functional context (Figure 6). At the protein-RNA interface (Figure 6A), pyrimidine-pyrimidine (YY) dinucleotides exhibit pronounced conformational preferences. The CC dinucleotide is significantly enriched in the canonical A-form conformer AA00 as well as the open conformer OP33. In contrast, UU preferentially adopts non-canonical and open conformations, including AB05, BB04, BB16, OP15, OP17 and OP26, while showing strong depletion of the canonical A-form AA00 state. The mixed pyrimidine step UC shows marked enrichment for the open conformers OP11 and OP33, whereas CU strongly prefers canonical and transitional conformers AA00 and AB06, respectively. Among pyrimidine-purine (YR) dinucleotides, UA displays significant enrichment in distorted and open conformations, including BB16, BB22, OP15, OP17, OP22 and OP32, while being underrepresented in canonical A-form conformers such as AA00, AA01 and AA07. The CG preferentially adopts AA00, AB07 and the Z-like conformer ZZ1S, but is depleted in AB05, BB04 and OP15, indicating strong sequence-dependent conformational selection. Within purine-pyrimidine (RY) dinucleotides, AU prefers AA11, OP11, OP21, OP25 and ZZ1S, suggesting enrichment of flexible and non-canonical geometries at the interface. Similarly, GU prefers AB05, BB04, OP21 and OP22, while showing reduced preference for widened and unwound A-form conformers AA01 and AA07. In contrast, GC strongly prefers the canonical A-form conformer AA00 and is significantly depleted in several non-canonical states, including AB05, BB04, BB16, OP15, ZZ1S and NANT. Among purine-purine (RR) dinucleotides, AA shows strong preference for transitional, intercalated and open conformers including AB06, BA16, BB22, IC09, OP15, OP21 and NANT, while being underrepresented in canonical A-form conformers AA00 and AA01. The GG dinucleotide prefers AA01 and the intercalated conformer IC08, whereas it is depleted in AB05, OP21 and NANT. Likewise, AG strongly prefers AA01, AA07 and ZZ1S, but shows negative preference for AA00 (Figure 6A).

**Figure 6.**
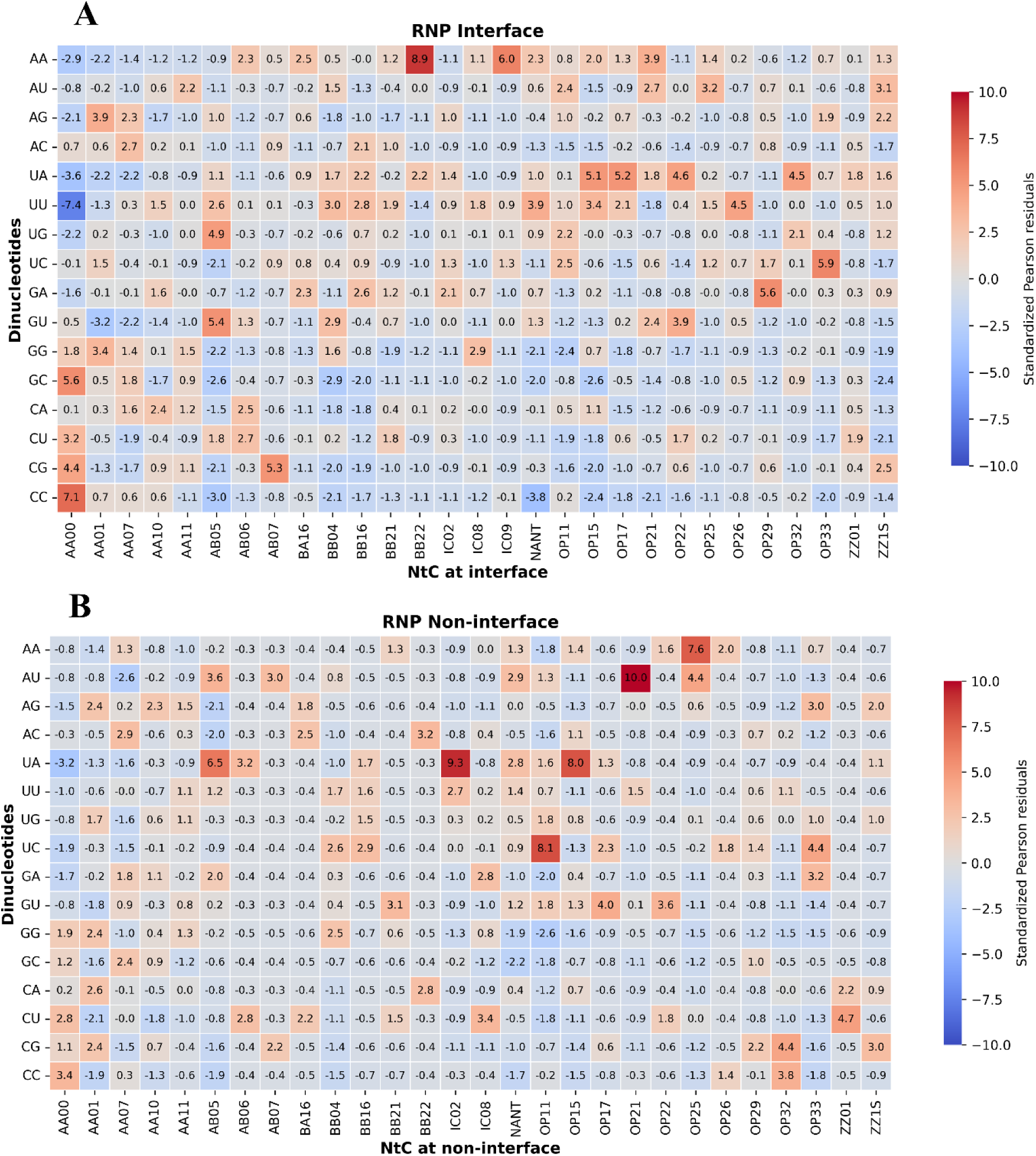
Standardized Pearson Residuals (SPR) of dinucleotide conformer preferences. SPR of dinucleotide conformers at the interface (A) and at the non-interface (B) are shown.

At the non-interface (Figure 6B), several distinct enrichment patterns are observed. Among YY dinucleotides, CC prefers AA00 and OP32, whereas UU prefers primarily the intercalated conformer IC02. The UC prefers BB04, BB16, OP11, OP17 and OP33, while CU prefers AA00, AB06, BA16, IC08 and ZZ01. The CG prefers AA01, AB07, OP29, OP32 and ZZ1S, indicating substantial conformational heterogeneity outside the interface. Among RY dinucleotides, AU prefers AB05, AB07, OP21, OP25 and NANT, whereas it is depleted in AA00. The GU prefers BB21, OP17 and OP22 conformers. For RR dinucleotides, AA shows enrichment predominantly in OP25, while AG prefers AA01, AA10, OP33 and ZZ1S, and is depleted in AB05 only. In contrast, GA prefers AB05, IC08 and OP33, while showing a negative preference for OP11. Similarly, GG prefers AA01 and BB04, but is depleted in OP11.

### Dinucleotide conformational preferences across protein-RNA structural classes

To investigate the relationship between RNA architecture and local backbone geometry, we have analyzed dinucleotide conformational preferences across a dataset divided into four protein-RNA structural classes defined in the Materials and Methods section. Distinct differences are observed between interface and non-interface regions, reflecting the influence of RNA structural context on conformational selection. The canonical A-form conformer AA00 emerged as the dominant NtC across all structural classes (Figure 7).

**Figure 7.**
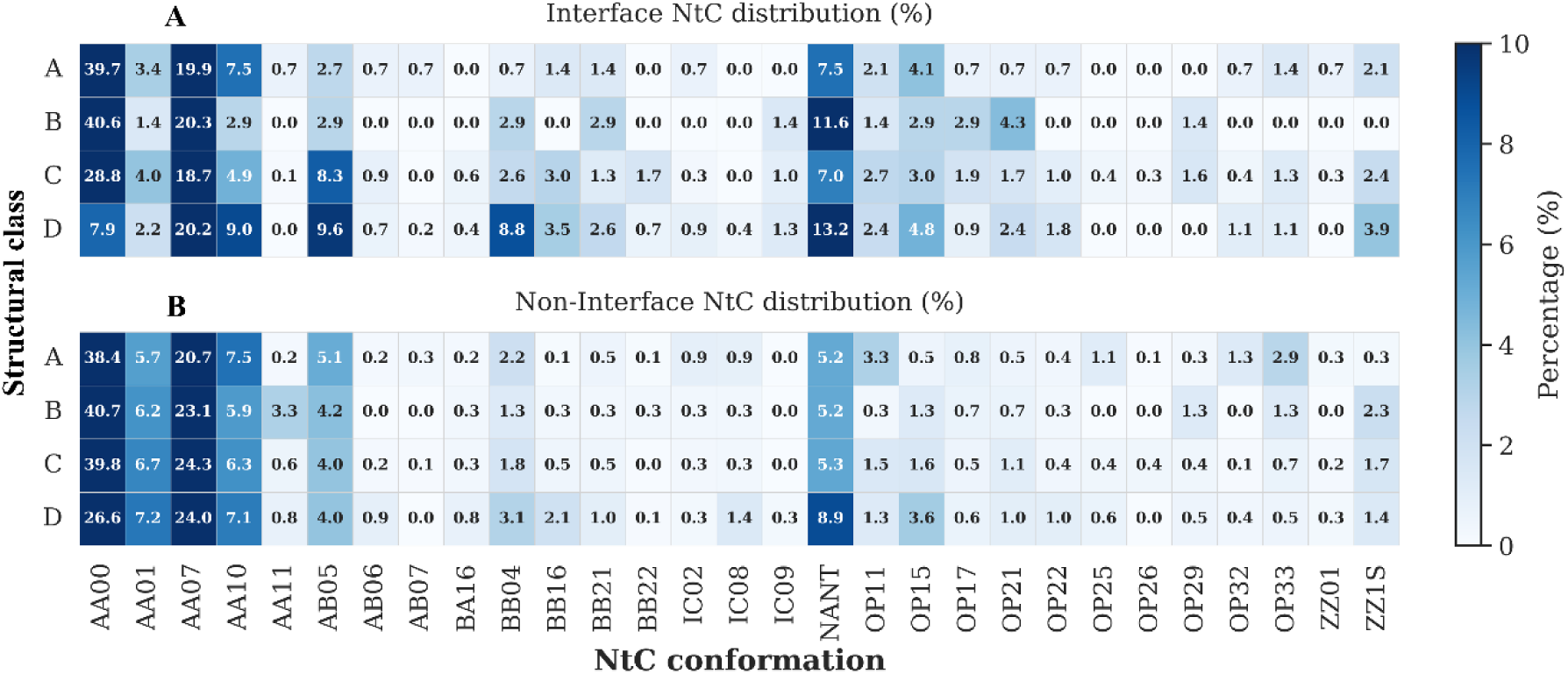
Percentage distribution of dinucleotide conformers (NtC) across four structural classes in the dataset of protein-RNA complexes: A (tRNA), B (rRNA), C (duplex RNA), and D (single-stranded RNA). (A) at the protein-RNA interface, (B) at the non-interface region.

At the protein-RNA interface (Figure 7A), AA00 is particularly enriched in Class A and Class B, accounting for 40.7% and 35.9% of all dinucleotide steps, respectively. At the non-interface regions (Figure 7B), AA00 is highly prevalent, especially in Class B, where it accounts for 50% of the conformers, and in Class C, where it nearly accounts for 41%. In contrast, AA00 is markedly underrepresented in Class D interfaces, where it accounts for only 13.9%, indicating reduced stabilization of canonical A-form geometry in flexible single-stranded regions. Similarly, AA00 shows comparatively lower abundance (36.5%) at the non-interface regions of Class A. In addition to AA00, the canonical A-like conformers AA01, AA07 and AA10 are consistently enriched across both interface and non-interface regions, reinforcing the predominance of A-form geometries in structured RNA environments. However, several non-canonical conformers display strong class-specific preferences. The A-B transitional conformer AB05 is particularly enriched in Class D interfaces (7.4%), suggesting increased backbone flexibility and conformational adaptability in single-stranded RNA. Interestingly, AB05 is more abundant at the non-interface regions of Class A, indicating that transitional geometries are also retained in structurally dynamic regions of folded RNAs such as tRNA. The distorted B-like conformers BB16, BB21 and BB22 are enriched at the protein-RNA interfaces across multiple structural classes compared to their corresponding non-interface regions. This enrichment suggests that local deviations from canonical A-form geometry are preferred during protein recognition. Overall, interface regions exhibit substantially greater conformational heterogeneity than non-interface regions, highlighting the dynamic structural adaptations associated with protein binding (Figure 7A). Distinct enrichment patterns are also observed for Z-like conformers. In Class A, the left-handed helical conformer ZZ01 is strongly preferred in both interface and non-interface regions, suggesting its potential role in stabilizing local tertiary architectures of tRNA. In contrast, the Z-like conformer ZZ1S is enriched at the interface in all structural classes, except Class A, indicating that left-handed or syn-associated geometries may be selectively stabilized during protein interaction. At the non-interface, however, ZZ1S was most abundant in Class B, suggesting structural context-dependent usage of Z-like conformations.

We further analyzed the proportion of dinucleotide steps that could not be assigned to any known NtC category (NANT). At the interface, NANT constitute a substantial fraction across all structural classes, accounting for 5.5% in Class A, 6.9% in both Classes B and C, and reaching 11.9% in Class D. The elevated occurrence of NANT conformers at interfaces suggests the presence of highly irregular or transient backbone geometries that may arise from local flexibility, induced-fit structural rearrangements or limitations in experimental resolution. In contrast, non-interface regions generally exhibit lower NANT frequencies, with 5.3% in Class A, 5.5% in Class B, 3.2% in Class C and 5.4% in Class D. The particularly low proportion observed in Class C indicates that canonical base-paired helices prefer to maintain well-defined and classifiable conformations away from protein-binding regions.

## Discussion

In this study, we systematically catalogue RNA dinucleotide conformers from a non-redundant dataset of 263 protein-RNA complexes using the standardized conformational framework proposed by Jiří Černý and colleagues (Černý et al. 2020). Historically, RNA structural analysis has been challenging due to fragmented classification schemes and inconsistent nomenclature, which limits comparative analyses and hinders a unified interpretation of RNA conformational space. The introduction of DNATCO framework provide a major advancement by establishing a comprehensive and standardized dinucleotide-level classification comprising 96+1 minor NtC classes and 15 broader CANA categories applicable to both DNA and RNA. Although DNATCO framework provides a robust global description of nucleic acid conformations, the original NtC definitions are derived from structurally diverse RNA molecules without explicitly considering their functional or interaction-specific contexts. Consequently, conformational states originating from distinct structural environments are grouped together within common clusters, potentially masking subtle but biologically meaningful variations associated with RNA recognition, folding and protein binding. Since RNA is inherently dynamic and highly responsive to its molecular environment, local conformational states observed at protein-binding interfaces may differ substantially from those present in canonical helical or non-interacting regions.

To address these limitations, we have refined RNA conformational states using a density-based clustering approach applied specifically to interface and non-interface regions of protein-RNA complexes. This strategy enabled the identification of conformational substates that are selectively enriched in protein-bound environments and revealed previously uncharacterized NtCs that are not represented in the existing classification system. Importantly, several newly identified conformers belong to transitional, open, intercalated and distorted B-like CANA classes, highlight the extensive structural adaptability of RNA during protein recognition. Our analyses demonstrate that protein-binding interfaces exhibit substantially greater conformational heterogeneity compared to non-interface regions. While canonical A-form conformers, such as AA00, remain dominant in structured RNA. Interface regions are enriched in conformations characterized by mixed sugar puckers, altered backbone torsions, disrupted base stacking and increased base separation. Such features are hallmarks of locally flexible or partially distorted RNA geometries and are consistent with previous observations that RNA undergoes conformational remodeling upon protein binding. These structural rearrangements facilitate mechanisms of induced fit and conformational selection (Murray et al. 2003b), enabling RNA molecules to optimize intermolecular contacts while preserving global structural integrity. Identification of novel conformers within the A-B transitional, OPN, ICL and miB classes further emphasizes that protein recognition frequently involves conformational states beyond canonical A-form helices. In particular, open and intercalated conformers observed at protein-binding interfaces promote accessibility of functional groups, accommodate steric constraints imposed by proteins and stabilize tertiary interactions required for complex formation. Similarly, the enrichment of distorted B-like and Z-like conformers suggests that local backbone plasticity contributes to sequence-specific recognition and structural adaptation (Placido et al. 2007; Butcher and Pyle 2011).

### Influence of B-factor RNA dinucleotide conformations

Crystallographic B-factors are widely used as indicators of local atomic mobility and conformational flexibility in biomolecular structures. Elevated B-factors often correspond to structurally dynamic or disordered regions, whereas lower values are associated with well-ordered conformations (Sun et al. 2019). Previous studies have demonstrated that B-factor profiles can effectively capture RNA flexibility and conformational heterogeneity, particularly in functionally important regions involved in molecular recognition and structural adaptation (Guruge et al. 2018; Keating et al. 2011). To evaluate the structural reliability and dynamic behavior of RNA conformations within the non-redundant set of 263 protein-RNA complexes, we analyzed crystallographic B-factors at both the structure and dinucleotide levels. Mean B-factor values and their corresponding standard deviations are calculated for each PDB structure and individual dinucleotide step, thereby capturing the average atomic displacement as well as local structural variability. The distribution of mean dinucleotide B-factors show a skewed pattern (Figure 8), with the majority of dinucleotide steps clustered below ∼60 Å². The overall distribution shows a mean B-factor of 51.1 Å² and a median value of 44.8 Å², indicating that most RNA conformations in the dataset occupy relatively low-to-moderate flexibility states, while a smaller subset corresponds to highly dynamic regions.

**Figure 8.**
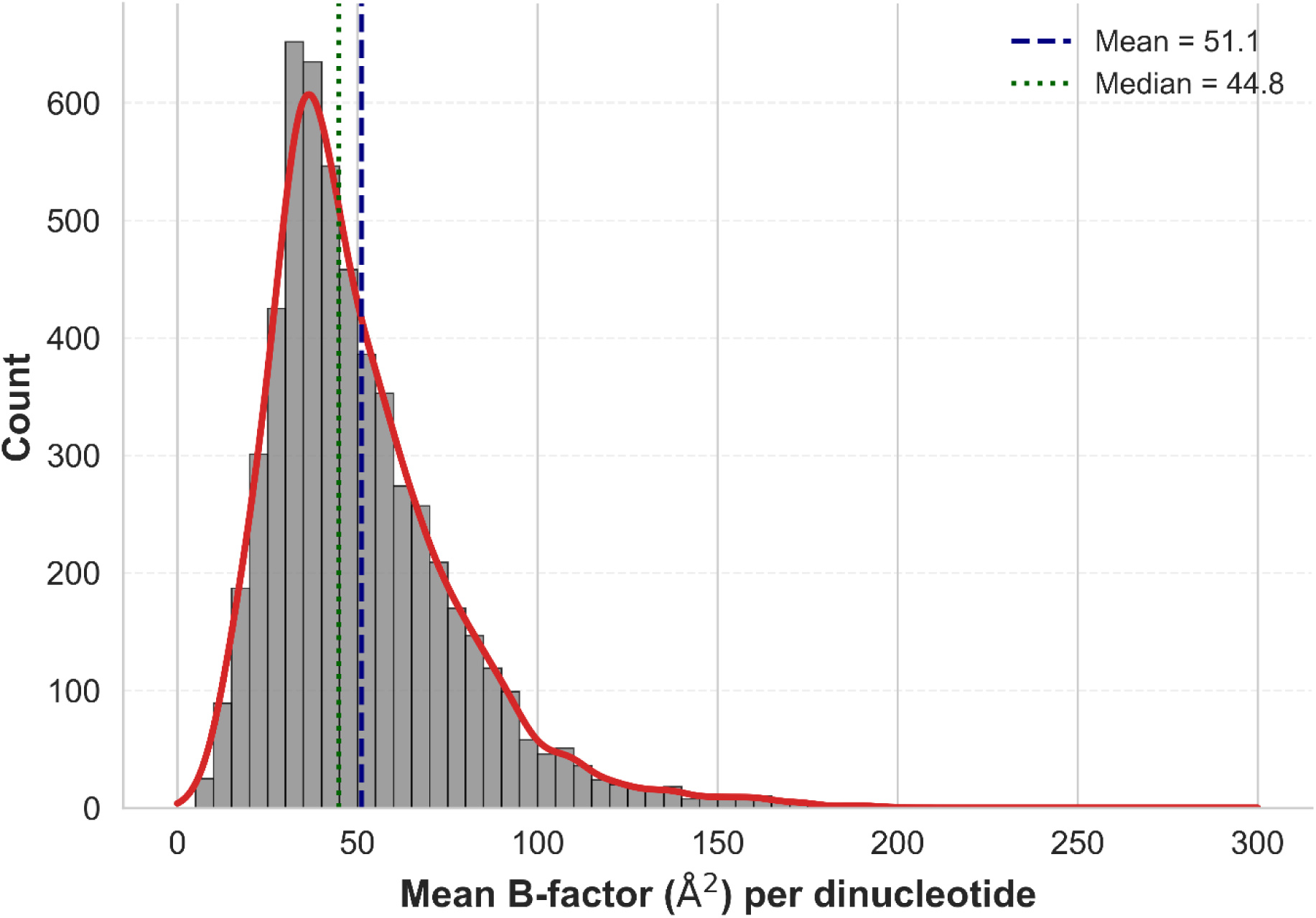
Distribution of RNA dinucleotides mean B-factors across protein-RNA complex dataset.

To further examine the relationship between structural flexibility and conformational reliability, we correlated the mean B-factor of each dinucleotide step with its corresponding Confal score for the newly identified NtC conformers (Supplementary Figure S4). The Confal score measures the degree of agreement between an observed dinucleotide geometry and its representative NtC cluster, with higher values indicating stronger conformational similarity and improved structural confidence. Most newly identified NtCs maintain a consistent high Confal score (>85) across a broad range of B-factor, indicating that these conformers remain structurally well-define despite local variations in atomic mobility.

Several conformers, including AB06, BB21 and OP32 (Figure S4A, C and E) exhibit weak positive correlations between B-factor and Confal score (*r* = 0.07–0.10), suggesting that moderate increases in local flexibility do not substantially affect conformational assignment quality. Similarly, BB22 (Figure S4D) show a negligible negative correlation (*r* = −0.061), indicating stable conformational behavior across different flexibility states. In contrast, some conformers exhibit moderate trends linking flexibility with conformational variability. The AB07 (Figure S4B) conformer show a moderate negative correlation (*r* = −0.563), indicating that higher B-factors are associated with reduced agreement with the representative cluster geometry. Likewise, the open conformer OP33 (Figure S4F) (*r* = −0.139) and the intercalated conformer IC08 (*r* = −0.194) display weak negative trends (Figure S4G), consistent with the intrinsically dynamic nature of open and non-helical RNA states, respectively. These conformers likely sample a broader conformational ensemble, resulting in greater structural heterogeneity with increased flexibility. Interestingly, the intercalated conformer IC09 (Figure S4H) exhibits a moderate positive correlation (*r* = 0.591), suggesting that structurally ordered instances of this conformer maintain strong similarity to the cluster representative despite variations in local mobility. Although several correlations are not statistically significant due to limited sample sizes, the overall trends provide important insights into the relationship between RNA flexibility and conformational stability. Our data show that most of the newly identified NtCs represent structurally robust conformational states rather than artifacts arising from highly flexible or poorly resolved regions. At the same time, the elevated B-factors observed in several open, transitional and intercalated conformers support the notion that local RNA flexibility contributes to conformational adaptability during protein recognition. These observations are consistent with previous studies demonstrating that crystallographic B-factors capture biologically relevant RNA dynamics and structural heterogeneity associated with molecular recognition and functional adaptation (Guruge et al. 2018; Sun et al. 2019).

### RNA conformational preferences in protein-RNA recognition

The diversity of RNA dinucleotide conformers identified in this study highlights the remarkable structural plasticity of RNA and its ability to adopt context-dependent backbone geometries influenced by sequence composition, base pairing and intermolecular interactions. The observed conformational repertoire spans canonical A-form helices, transitional A/B geometries, distorted B-like states, open conformations and compact intercalated structures with unusual torsional arrangements. Such diversity underscores the dynamic nature of RNA and suggests that local backbone flexibility plays a central role in facilitating protein recognition and stabilizing protein-RNA complexes. Among the newly identified conformers, AB06 and AB07 NtCs belong to A-B transitional CANA class and represent intermediate geometries between canonical A-form and B-form conformations. These conformers are predominantly enriched at protein-binding interfaces, indicating their functional relevance in molecular recognition. In particular, AB06 frequently participates in intermolecular H-bonds. A representative example is observed in the *A. aeolicus* NusB-NusE ternary complex bound to BoxA RNA (PDB ID: 3R2C), which functions in transcription antitermination. In this complex, G2-C3 dinucleotide adopts an AB06-like geometry that facilitates specific recognition by surface residues of the NusB protein (Stagno et al. 2011). Such transitional conformations may therefore provide the geometric adaptability required for optimizing intermolecular contacts during RNA recognition. Similarly, the distorted B-like conformers BB21 and BB22, classified within the minor B-like (miB) CANA class, are preferentially enriched at protein-binding sites. These conformers exhibit substantial backbone rearrangements while retaining partial B-form characteristics, suggesting that local distortion of canonical RNA geometry may contribute to interface formation. The functional relevance of BB21-like conformation is exemplified by the *S. cerevisiae* She2p-ASH1 mRNA localization complex (PDB ID: 5M0I), where C27-G28 dinucleotide undergoes pronounced conformational rearrangement upon protein binding, resulting in a kinked RNA architecture that promotes recognition by the She2p protein (Edelmann et al. 2017). These observations indicate that distorted B-like geometries may serve as adaptable structural motifs that accommodate protein-induced RNA deformation.

The newly identified RNA open conformers, OP32 and OP33, belong to the OPN CANA class and are characterized by reduced base stacking, increased base separation and enhanced backbone flexibility. These conformers represent structurally dynamic RNA states that deviate from canonical helical geometries and are frequently associated with functionally important flexible regions. OP32 and OP33 are predominantly observed in tRNA loop regions, where local structural adaptability is required for tertiary interactions and molecular recognition. Their enrichment in protein-bound environments suggests that open conformations may facilitate conformational rearrangements necessary for protein recognition, induced fit and stabilization of ribonucleoprotein complexes (Butcher and Pyle 2011; Das et al. 2023). The newly identified intercalated conformers IC08 and IC09, assigned to the ICL CANA class, represent compact yet non-helical RNA states characterized by mixed B-A sugar puckers and pronounced backbone distortions. Both conformers maintain anti-glycosidic orientations and elevated μ torsions, whereas IC09 additionally exhibits a characteristic ζ/α torsional switch. Notably, IC09 is observed almost exclusively at protein-RNA interfaces, where it contributes to stabilizing intermolecular interactions, particularly through intermolecular H-bonds. A representative example is found in *S. cerevisiae* Nab2p-poly(A) RNA complex (PDB ID: 5L2L), in which H-bond between N6 of A11 and sulfur atom of a cysteine residue promotes Zn-finger dimerization and stabilizes RNA recognition through a unique spatial arrangement of zinc-finger domains (Aibara et al. 2017).

The enrichment of IC09-like conformers at the interfaces suggests that compact intercalated geometries may facilitate highly specific recognition events within ribonucleoprotein assemblies. Collectively, these findings demonstrate that protein-RNA recognition is frequently associated with non-canonical and transitional RNA conformational states rather than exclusively canonical A-form helices. The enrichment of open, distorted, transitional and intercalated conformers at protein-binding interfaces supports the idea that RNA undergoes localized conformational adaptation to optimize intermolecular interactions. Such structural flexibility likely contributes to mechanisms of induced fit and conformational selection during ribonucleoprotein assembly.

## Conclusion

In this study, we have systematically analyzed RNA dinucleotide conformers to investigate the structural variability of RNA and its role in protein-RNA recognition. By applying advanced density-based clustering approach to the high-dimensional torsional space of RNA dinucleotide steps, we identify local conformational states independent of global RNA architecture. Our analyses demonstrate the predominance of the canonical A-form conformer AA00 across the diverse RNA structural contexts, while also revealing substantial enrichment of non-canonical and transitional conformers at protein-binding interfaces. Notably, we identify several previously unreported NtCs, including A-B transitional conformers AB06, AB07, BB21, BB22, OP32, OP33, IC08 and IC09. These are preferentially associated with protein-interacting regions and likely contribute to RNA conformational adaptation during molecular recognition. The observed enrichment of open, intercalated and distorted conformers at protein-RNA interfaces highlights the importance of local backbone flexibility in facilitating intermolecular interactions. A significant fraction of dinucleotide steps remains unassigned to existing NtC classes, particularly at protein-binding interfaces, suggesting the presence of rare, transient or highly context-dependent RNA conformations that are not fully represented in current classification schemes. This may partly reflect the limited availability of high-resolution RNA structural data and challenges associated with refinement of flexible RNA regions in experimentally determined structures. Importantly, this work introduces a refined RNA dinucleotide conformer library derived specifically from protein-bound RNA environments. The library captures sequence-dependent and position-specific conformational preferences, including newly identified motifs associated with tRNA structural elements and stacking-mediated interactions. By linking RNA sequence, local geometry and intermolecular recognition, this conformer library provides an improved structural framework for studying RNA flexibility and conformational dynamics in ribonucleoprotein complexes. Collectively, our findings expand the current understanding of RNA conformational space and provide valuable resources for RNA structural modeling, docking, molecular dynamics simulations and the development of next-generation computational approaches for predicting protein-RNA interactions. The complete RNA NtC conformer library developed here is available at https://github.com/shrikantcombio/RBPs_RNA_conformers_lib.git.

## Acknowledgments

SK and SM are thankful to IIT Kharagpur for providing the fellowship and infrastructure facility. RPB acknowledges the Department of Biotechnology (DBT), Govt. of India (grant no. BT/PR40175/BTIS/137/41/2022) for the Bioinformatics Centre.

## Authors contributions

S.K. conceptualized the work and wrote the initial draft. S.K. and S.M. derived the RNA conformers library, formal analysis, visualization, and edited the manuscript. R.P.B. conceptualized the work, analyzed data, edited the final version of the manuscript, overall supervised the project, and provided the resources.

## Data availability

The library developed in this study is available on GitHub at https://github.com/shrikantcombio/RBPs_RNA_conformers_lib.git.

