## Supplementary Material for "Deciphering conformational preferences of RNA in protein-RNA recognition"

**Supplemntary material for:**

*Corresponding author

Ph: +91-3222-283790

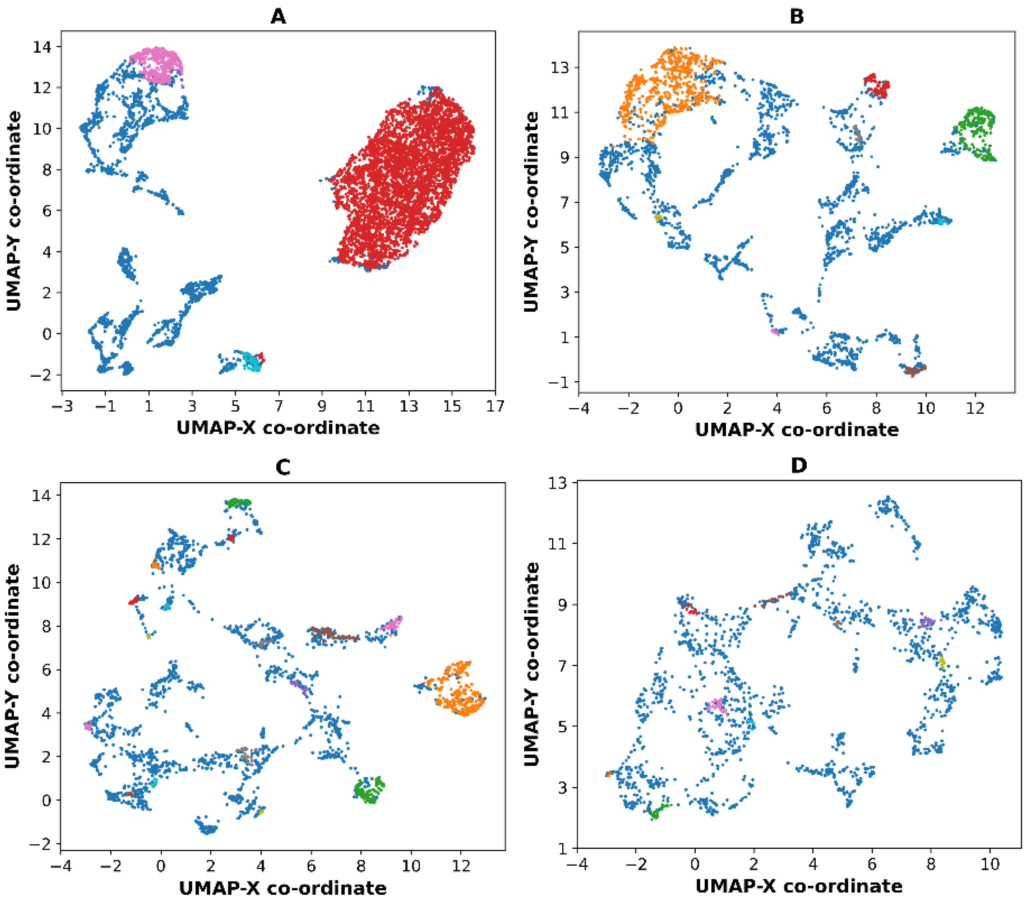


Figure S1. UMAP projection of each of the four iterations including outlier after clustering using DBSCAN. We find 29 clusters, each representing a group of highly similar conformers. The largest cluster is presented with red colour in first iteration plot.

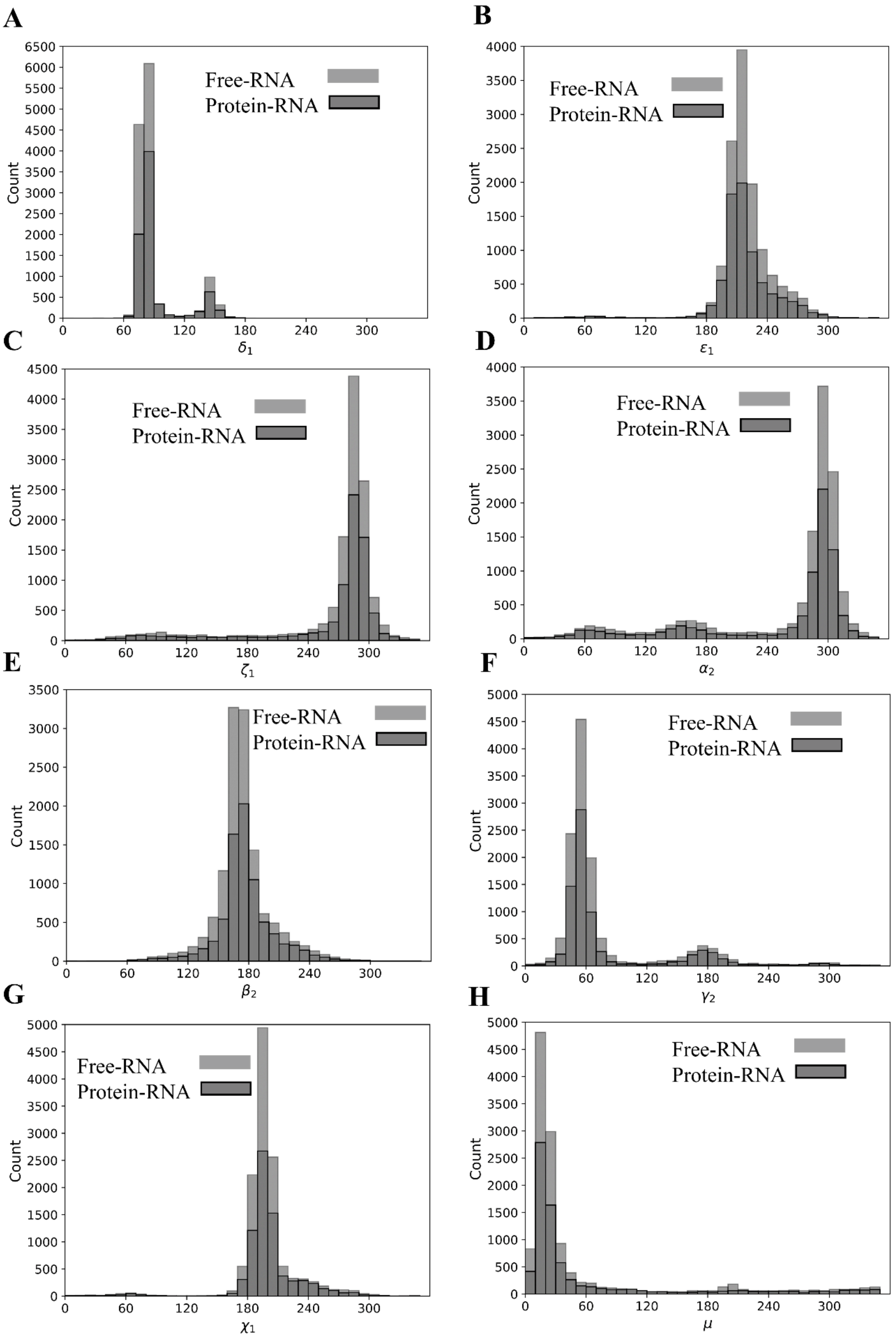


Figure S2. Distributions of the backbone dihedral angles of each nucleotide present in protein-RNA complexes and in free RNA structures.

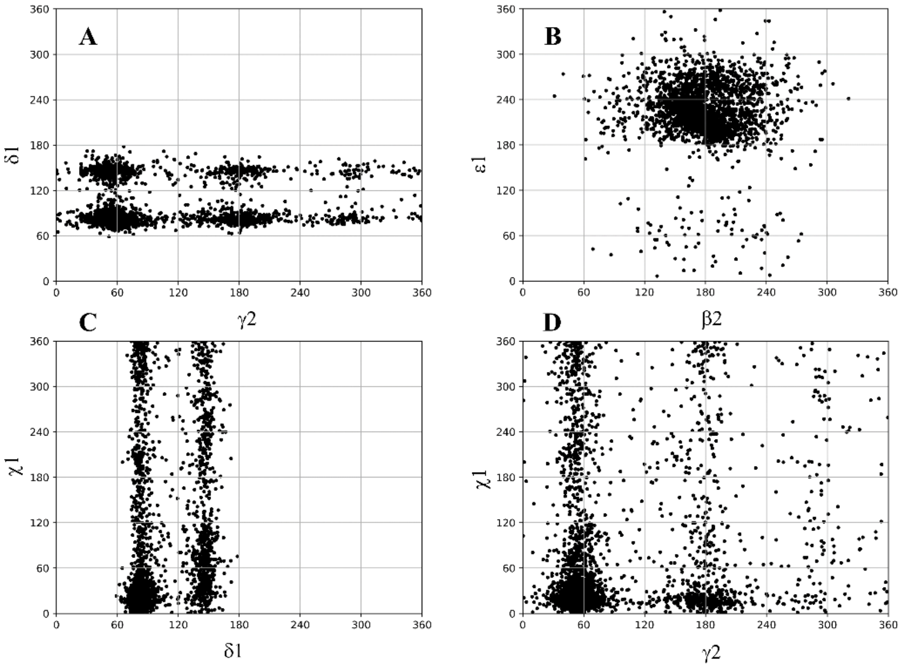


Figure S3. Two-dimensional scattergrams illustrating the distribution and correlation between various backbone and glycosidic torsion angles in 288 protein-RNA complexes. (A) between $\gamma_2$ and $\delta_1$, (B) between $\beta_2$ and $\varepsilon_1$, (C) between $\delta_1$ and $\chi_1$ and (D) between $\gamma_2$ and $\chi_1$.

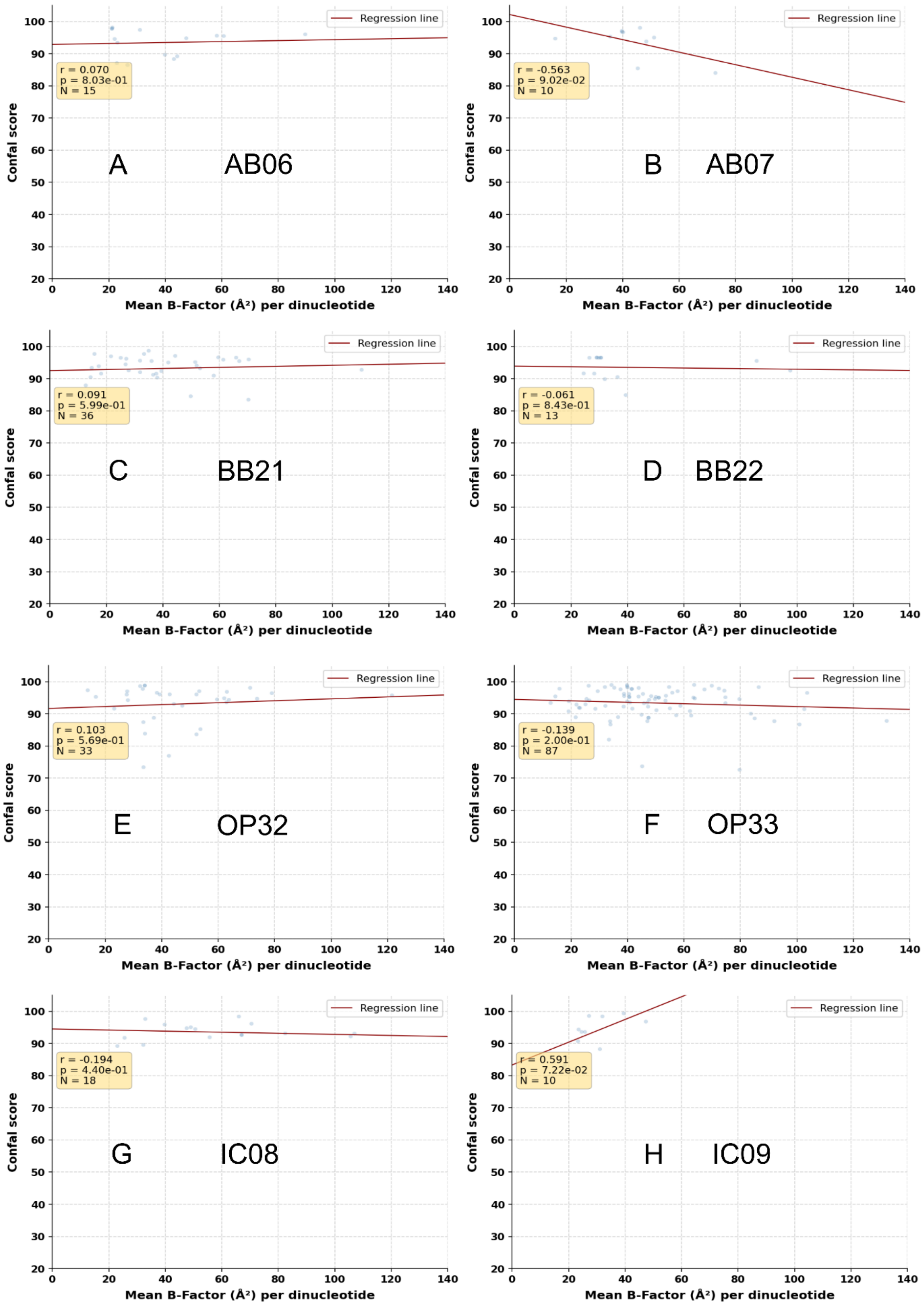


Figure S4. Relationship between structural flexibility and dinucleotide conformation quality of newly identified NtCs: Correlation analysis between the mean B-factor per dinucleotide and the corresponding Confal score for the newly identified NtCs: (A) AB06, (B) AB07, (C) BB21, (D) BB22, (E) OP32, (F) OP33, (G) IC08, and (H) IC09.

Supplementary tables are provided in Supplementary_Tables.xlsx.
